# The XenCart Protocol: A Method for Alcian Blue Labeling and Quantitative Analysis of Craniofacial Cartilage in *Xenopus*

**DOI:** 10.64898/2026.05.30.728963

**Authors:** Umar Aziz, Leona Bhandari, Claudia Lizama, Riya Maurya, Amanda J.G. Dickinson

## Abstract

Craniofacial birth defects, such as cleft lip and palate, are among the most common congenital anomalies and often arise from disruptions in early facial patterning. Many of these defects are linked to environmental teratogens, yet such exposures cannot be directly tested in humans, making animal models essential for evaluating developmental risks. *Xenopus laevis* offers a powerful solution: its tadpoles develop externally, share deeply conserved craniofacial patterning mechanisms with humans, and provide an accessible platform for uncovering how environmental exposures reshape facial structures during development. Here, we present the XenCart Protocol, a reproducible workflow for Alcian Blue staining and quantitative morphometric analysis of Xenopus craniofacial cartilage. This method provides clear visualization of individual cartilage elements and can be readily applied to investigate genetic or environmental perturbations. The Xenopus craniofacial skeleton contains distinct cartilaginous structures that perform key biomechanical functions and share strong homology with regions of the human craniofacial skeleton. These similarities allow direct comparison of developmental outcomes across vertebrates. As part of a CURE-based undergraduate course, the XenCart Protocol was used to measure jaw cartilage dimensions in tadpoles exposed to an emerging teratogen, e-liquids used in vaping. E-liquid exposure caused consistent reductions across major craniofacial cartilages, including shorter Meckel’s cartilage, narrowed infrarostral width, decreased basihyobranchial and ceratohyal dimensions, and reduced suprarostral angles, reflecting an overall shift toward a smaller, more compact craniofacial morphology. These patterns suggest potential disruption of neural crest cell migration or signaling pathways for craniofacial cartilage development, mechanisms that, if similarly affected in humans, could contribute to midfacial narrowing, jaw underdevelopment, or increased vulnerability to conditions such as orofacial clefts. The ability to detect robust, structure-specific differences highlights the sensitivity of the protocol and its strong alignment with student-led research. These findings also pinpoint the precise regions of the jaw most affected by e-liquid exposure, providing a foundation for uncovering the developmental mechanisms driving these craniofacial changes. In summary, the XenCart Protocol provides a standardized, scalable method for quantifying craniofacial cartilage development and offers a powerful platform for both mechanistic research and undergraduate training in developmental biology and toxicology.

## Introduction

Craniofacial birth defects, including cleft lip and palate, are among the most common congenital anomalies and present major clinical, developmental, and societal challenges (Wehby and Cassell 2010; Vyas et al. 2020; Salari et al. 2022). These abnormalities can arise from both genetic perturbations and environmental teratogens that disrupt craniofacial patterning (Dixon et al. 2011). Animal models are essential for understanding the causal and mechanistic factors underlying craniofacial defects in ways that cannot be directly tested or fully interpreted in humans (Van Otterloo et al. 2016; Ibarra and Atit 2020). One such model, Xenopus, provides a particularly powerful system for studying craniofacial development due to their external development, optical accessibility, and highly conserved craniofacial development (Dickinson 2016; Dubey and Saint-Jeannet 2017; Kong et al. 2025). In addition to supporting basic and translational research, Xenopus-based assays are well suited for independent and Course-based Undergraduate Research Experiences (CUREs)(Olive et al. 2003), where students can engage in authentic experimental design, hypothesis testing, and quantitative analysis of craniofacial phenotypes resulting from defined perturbations.

Craniofacial development can be assessed directly by examining the skeletal elements that give the head its shape, structural support, and functional organization. In *Xenopus* tadpoles, the craniofacial cartilages also provide a particularly informative readout of overall craniofacial morphology. The craniofacial skeleton of *Xenopus* tadpoles consists of a series of distinct cartilaginous elements that support feeding, respiration, and jaw movement (summarized in Fig 1A)(Lukas and Olsson 2018; MacKenzie et al. 2022). These structures mirror key regions of the human craniofacial skeleton, enabling comparison of developmental processes across vertebrates (Palpant et al. 2015; Dubey and Saint-Jeannet 2017). In amphibians the suprarostral plate, palatoquadrate, and ethmoid/trabecular region form a continuous dorsal craniofacial cartilage complex. The suprarostral plate is the anterior upper-face portion, the palatoquadrate is the paired lateral upper-jaw support, and the ethmoid region is the anterior midline neurocranial/nasal scaffold. Although these elements have different developmental and anatomical identities, they are physically integrated and together support the upper face and roof/lateral margins of the mouth (MacKenzie et al. 2022). The ventral portion of the tadpole jaw consists of the anterior most infrarostral cartilage which forms the oral rim of the mouth and the paired Meckel’s cartilages which are the major lower jaw support (MacKenzie et al. 2022). In mammals the Meckel’s cartilage serves as a scaffold for mandibular development before contributing to the malleus and incus (Anthwal et al. 2017). Importantly, the palatoquadrate of the upper jaw articulates with the Meckel’s cartilage of the lower jaw to establish the primary jaw joint (Anthwal and Tucker 2023). Posterior to the jaw structures are the paired ceratohyal cartilages and a single midline basihyobranchial cartilage which stabilize the floor of the mouth and supports pharyngeal movements(MacKenzie et al. 2022). Posterior to basihyobranchial are the ceratobranchials that form the branchial basket, providing structural support for feeding and respiratory function during larval stages (MacKenzie et al. 2022; Rose 2023). Together, the ceratohyal, basihyobranchial, and ceratobranchials are homologous to elements of the mammalian hyoid and pharyngeal apparatus, which support swallowing, speech, and airway stability (Gillis et al. 2013; Zhou et al. 2019; Lukas and Ziermann 2022). Collectively, these cartilages underscore the close evolutionary and functional relationship between amphibian and mammalian craniofacial architecture.

**Figure 1.**
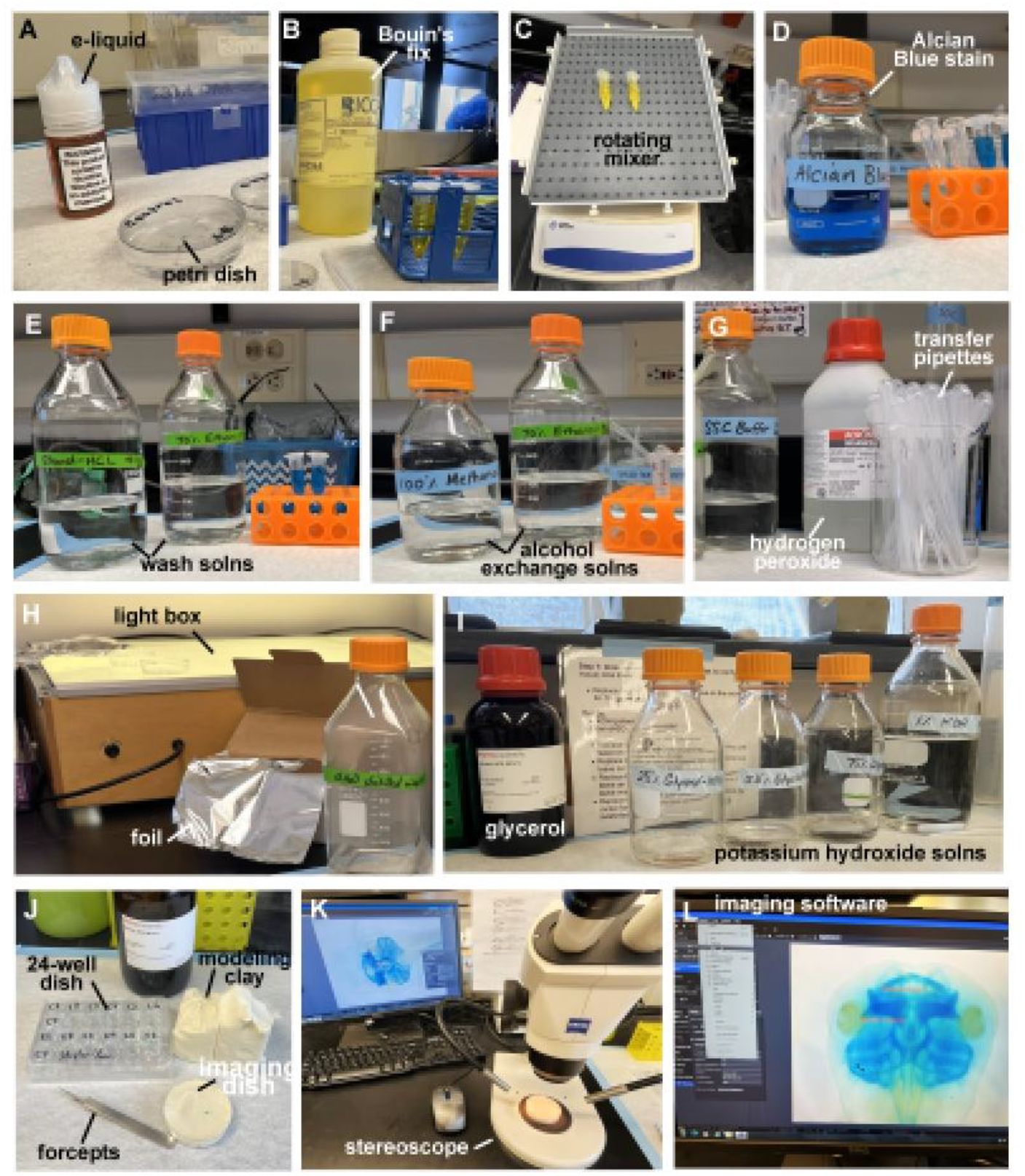
Reagents, supplies, and equipment used for craniofacial cartilage staining and imaging. Representative images of the materials used to process, stain, clear, and image *Xenopus laevis* embryos for craniofacial cartilage analysis. (A) E-liquid treatment reagent and petri dish used for embryo exposure. (B) Bouin’s fixative used for specimen fixation. (C) Rotating mixer used to incubate embryos during fixation, staining, washes, and solution exchanges. (D) Alcian Blue stain used to label craniofacial cartilage. (E) Wash solutions used to remove excess stain and prepare specimens for clearing. (F) Alcohol exchange solutions used during dehydration and processing. (G) Hydrogen peroxide and transfer pipettes used for bleaching and specimen handling. (H) Light box and, foil used during depigmentation steps. (I) Glycerol and potassium hydroxide solutions used for tissue clearing. (J) Imaging supplies, including 24-well dishes, modeling clay, imaging dishes, and forceps, used to position stained embryos. (K) Stereoscope and computer setup used to image the stained craniofacial cartilages. (L) Software used to measure stained specimens.

Here, we describe a protocol for labeling craniofacial cartilages with Alcian Blue and measuring individual cartilage elements that we call the **XenCart Protocol**. This protocol provides clear visualization of individual cartilages, enables precise morphometric measurements, and is adaptable for both research and undergraduate teaching laboratories. Due to its simplicity this method also offers a robust and scalable platform to investigate how genetic disruption or environmental insults alter craniofacial development.

As a brief demonstration of the method’s utility, we apply the XenCart protocol to tadpoles exposed to a chemical we have found to act as a teratogen in *Xenopus* embryos to illustrate its sensitivity in detecting region-specific craniofacial cartilage defects. Our previous work showed that vanillin-flavored e-liquids can produce striking craniofacial abnormalities in Xenopus tadpoles, including midfacial narrowing and oral clefts, potentially through disruption of retinoic acid signaling (Kennedy et al. 2017; Dickinson et al. 2021). Other studies similarly suggest that components of e-liquids can interfere with developmental processes raising concerns about their toxicity (Palpant et al. 2015; Shi et al. 2019; Onyenwoke et al. 2022; Ozekin et al. 2023; Black et al. 2025). These findings underscore the need for a standardized, element-specific approach capable of quantifying how craniofacial cartilages respond to environmental exposures.

In conclusion, the broad goal of this protocol is to provide an accessible and flexible approach for analyzing the craniofacial skeleton under a wide range of genetic or environmental perturbations. This workflow offers a simple quantitative platform for developmental biology, toxicology, and CURE-based undergraduate research.

## Materials and Methods: The XenCart Protocol

### A. Preparing Reagents

1. **Ethanol (100%)**: Use absolute (100%) ethanol as provided, without dilution.
2. **Alcian Blue Staining Solution**: Make an Alcian Blue solution by first making an ethanol and acetic acid solution (mix 400 mL of 100% ethanol with 100 mL of acetic acid). Then dissolve 0.05-0.1g Alcian Blue powder into the acid-ethanol solution and mix overnight. Next filter the solution using coarse filter paper to remove any undissolved particles. Note that the solution will look too dilute just after mixing but will darken by the next day. The ideal amount of Alcian Blue powder can vary depending on its age and possibly other factors, testing it on wildtype embryos is recommended before starting your experiment.
3. **70% Ethanol**: Make 70% ethanol by diluting 100% ethanol with distilled water to a final concentration of 70% (v/v).
4. **Ethanol + Hydrochloric Acid (HCl) Solution**: Make an ethanol/HCl solution by mixing 5 mL of 1 N HCl with 495 mL of 70% ethanol.
5. **70% Methanol**: Make 70% methanol by diluting 100% methanol with distilled water to a final concentration of 70% (v/v).
6. **50% Methanol**: Make 50% methanol by diluting 100% methanol with distilled water to a final concentration of 50% (v/v).
7. **25% Methanol**: Make 25% methanol by diluting 100% methanol with distilled water to a final concentration of 25% (v/v).
8. **Phosphate Buffered Saline with Tween (PBT)**: Make PBT by dissolving 8 g NaCl, 0.2 g KCl, 1.44 g Na_2_HPO_4_, and 0.24 g KH_2_PO_4_ in 800 mL distilled water. Adjust the pH to 7.4 and then add 2 mL Tween-20, and bring the final volume to 1 L with distilled water.
9. **20× Saline–Sodium Citrate (20× SSC)**: Make 20× SSC by dissolving 175.3 g NaCl and 88.2 g sodium citrate dihydrate in distilled water, adjusting the pH to 7.0, and bringing the final volume to 1 L.
10. **2× Saline–Sodium Citrate (2× SSC)**: Make 2× SSC by diluting 100 mL of 20× SSC with distilled water to a final volume of 1 L.
11. **Depigmentation Solution (Prepare Fresh)**: Make the depigmentation solution by mixing 9 mL of 2× SSC with 1 mL of 50% hydrogen peroxide. Add 1 mL formamide and 9 mL distilled water immediately before use.
12. **1% Potassium Hydroxide (KOH)**: Make a 1% KOH solution by dissolving 10 g potassium hydroxide in 1,000 mL distilled water.
13. **25% Glycerol KOH**: Make 25% glycerol KOH by diluting 100% glycerol to 25% (v/v) using 1% KOH as the diluent.
14. **50% Glycerol KOH**: Make 50% glycerol KOH by diluting 100% glycerol to 50% (v/v) using 1% KOH as the diluent.
15. **75% Glycerol KOH**: Make 75% glycerol KOH by diluting 100% glycerol to 75% (v/v) using 1% KOH as the diluent.
16. **Bouin’s Fixative** (purchase already prepared)
17. **Buffered Tricaine solution**. Prepare a 2.5% (w/v) tricaine (MS-222) stock by dissolving 250 mg MS-222 in water to a final volume of 10 mL, then adjust the solution to pH 7.0–7.5 with Tris (and store cold, protected from light).

### B. Required Tools and Equipment

1. White modeling clay
2. Any size plastic petri dishes
3. Semi-fine tip Tweezers
4. Disposable pipettes
5. 24, 48 or 96 well plate for storing individual tadpoles.
6. Modify a disposable pipette by cutting off the tip to create a wider opening for transferring tadpoles
7. Imaging Tray: Modify a 5cm plastic petri dish by pressing into half of the base with the white modeling clay, flattening and smoothing the surface, then adding a thin layer of 100% glycerol.
8. Benchtop rotator or shaker

Reagents, tools and equipment are summarized in Figure 1 and Table 1.

**Table 1:** Materials and Reagents.

| REAGENT/MATERIAL | CATALOG # | COMPANY |
| --- | --- | --- |
| Ethanol, absolute (100%) | V1101 | Koptec |
| Methanol, absolute (100%) | A452-4 | Fisher Chemical (Fisher Scientific) |
| Alcian Blue 8GX powder | 400450100 | Acros organics |
| Acetic acid (glacial) | A35-500 | Fisher Scientific |
| Hydrochloric acid, 1 N | W0603D/G | Titristar |
| Potassium hydroxide (KOH) | 124060010 | Acros organics |
| Sodium chloride (NaCl) | BP358-212 | Fisher Scientific |
| Potassium chloride (KCl) | BP366-500 | Fisher Scientific |
| Disodium phosphate ($\text{Na}_2\text{HPO}_4$) | S-0876 | Millipore Sigma |
| Monopotassium phosphate ($\text{KH}_2\text{PO}_4$) | P-284 | Fisher Scientific |
| Tween-20 | BP337-100 | Fisher Scientific |
| Sodium citrate dihydrate | W302600 | Millipore Sigma |
| Hydrogen peroxide (50%) | 302865000 | Acros Organics |
| Formamide | F7503 | Millipore Sigma |
| Glycerol (100%) | BP229-1 | Fisher Scientific |
| Disposable transfer pipettes | 13-711-7M | Fisher Scientific |
| 15 mL conical tubes | 28-103 | Genesee Scientific |
| 5 cm plastic petri dishes | 25384-090 | VWR International |
| 24-well culture plates | TR5002 | TrueLine |
| tricaine (MS222) | TRs1 | Western Chemical Inc. |
| Bouin's Fixative | HT10132 | Millipore Sigma |
| 5ml microcentrifuge tubes | 50-005-49282 | Fisher Scientific |
| Aluminum foil, 1 roll | local grocery store |  |
| White modeling clay, 1 pack | local art store |  |

### C. Embryo Collection and Fixation

1. **Collect Mutant, Morphant or Teratogen exposed embryos** at developmental stages 45-46 (older stages can also be used).
2. **Anesthesia**. To anesthetize embryos at 0.01% of tricaine (MS-22 2), add 40 µL of 2.5% tricaine stock directly to 10 mL of frog embryo buffer containing the embryos and gently mix by swirling. Anesthetize embryos for 10 minutes or until they no longer respond to touch.
3. **Fixation**. Transfer anesthetized embryos to a centrifuge tube, remove the buffer, and replace it with Bouin’s fixative. Place on rotator or shaker overnight at room temperature and then place in fridge for up to 6 months. Other fixatives such as 4% paraformaldehyde may offer similar results.

### D. Alcian Blue Staining

We present here a flexible staining procedure to visualize cranial cartilage based on published protocols (Kelly and Bryden 1983; Walker and Kimmel 2007; Di Renzo et al. 2011).

1. **Remove fixative and replace it with ethanol**. This step is to remove the fixative.
  1.1) Remove Bouin’s fixative from the tube containing fixed tadpoles using a plastic transfer pipette
  1.2) Add 70% ethanol to the top of the tube.
  1.3) Repeat 70% ethanol washes several times over a period of 3 days ensuring the fluid is no longer yellow. On the second wash day change the tubes since some yellow will be stuck to the tubes.
2. **Alcian Blue staining** This step stains cartilage so craniofacial structures can be visualized.
  2.1) Remove the ethanol from the tube containing fixed tadpoles.
  2.2) Fill the tube completely with Alcian Blue staining and place on shaker or rotator.
  2.3) Incubate tadpoles in this staining solution for 2 days at room temperature to allow the dye to penetrate cartilage tissue. The incubation time can be flexible from 2-3 days.
3. **Removal of excess stain** These washes remove unbound dye and reduce background staining.
  3.1) Remove the Alcian Blue solution and replace it with 70% ethanol.
  3.2) Place the tube on a rocker for 10 minutes.
  3.3) Replace with fresh 70% ethanol and repeat rocking two more times for a total of three washes (30 minutes total). This step is also very flexible and each exchange can be extended to 1 hour.
4. **Ethanol–HCl Wash** This step further removes background staining and improves image clarity.
  4.1) Replace the 70% ethanol with ethanol plus hydrochloric acid and rock for 10 minutes.
  4.2 Replace with fresh ethanol plus hydrochloric acid solution and rock for an additional 30 minutes. These steps are also flexible within 1 hour.
5. **Alcohol Exchange (Ethanol to Methanol)** This step helps to prevent excessive bubbles during the depigmentation step possibly by the permeabilization of the skin.
  5.1) Replace the ethanol–HCl solution with 70% ethanol and rock for 10 minutes to remove acid.
  5.2) Replace ethanol with 70% methanol and rock for 30 minutes to exchange alcohols.
  5.3) Repeat by replacing with new 70% methanol and rock for again overnight. The last step is very flexible and tadpoles may remain in 70% methanol for up to 3 days but keep in the fridge.
6. **Buffer Exchange** This step gradually replaces methanol with a buffered solution.
  6.1) Replace 70% methanol with 50% methanol and rock for 10 minutes.
  6.2) Replace 50% methanol with 25% methanol and rock for 10 minutes.
  6.3) Replace 25% methanol with PBT buffer and rock for 30 minutes.
  6.4) Replace the PBT with new PBT. At this point the embryos can be placed in the fridge overnight or proceed to the next step.
7. **Pigment Removal (Depigmentation)** This step removes natural pigmentation so cartilage structures are easier to see.
  7.1) Transfer tadpoles using a cut transfer pipette to a small petri dish, and remove the PBT while holding the dish on an angle.
  7.2) Add freshly prepared depigmentation solution, place on a light box, and loosely cover with foil to maximize light exposure.
  7.3) Monitor tadpoles until dark pigment is removed and eyes appear light yellow. The time this takes can vary from 20 minutes to 2 hours and should be closely monitored to ensure that tadpoles do not get too white which can result in increased fragility and poorer imaging results.
  7.4) Remove the depigmentation solution and replace it with PBT buffer.
  7.5) Transfer tadpoles using a cut transfer pipette to a clean 1-5 mL tube and rock for 5 minutes.
  7.6) Replace the PBT buffer 3 times, rocking for 5 minutes between each change. Note: Tadpoles may be stored in buffer at 4°C for 1-4 days.
8. **Tissue Clearing** This step makes tissues transparent, so internal cartilage structures are visible.
  8.1) Replace PBT buffer with 1% potassium hydroxide (KOH) and rock for 1 hour.
  8.2) Replace with 25% glycerol in 1% KOH and rock for 30 minutes-1hour.
  8.3) Replace with 50% glycerol in 1% KOH and rock for 30 minutes-1hour.
  8.4) Replace with 75% glycerol in 1% potassium hydroxide and rock for 30 minutes-1hour
  8.5) Replace with 100% glycerol and incubate for at least 24 hours. Samples may be stored indefinitely in glycerol.

### E. Imaging Craniofacial Cartilage

This step captures standardized images for comparison and measurement.

1. Add 100% glycerol to a clay-lined petri dish, using just enough to cover the tadpole.
2. Transfer one tadpole at a time into the dish.
3. Use forceps to position the tadpole in the center of the dish with the head facing upward in a ventral view. A depression can be made in the clay to improve the view and keep the tadpole from rolling or drifting in the dish.
4. Photograph the head using a high-resolution light microscope with a digital camera at a constant magnification.
5. Keep magnification, lighting, exposure, and contrast consistent for all images and avoid strong shadows.
6. Save all images as software specific files as well as .tif or jpeg files for image compilation.
7. After imaging, transfer each tadpole to a labeled well of a 24, 48 or 96-well plate. If necessary, the embryo can be reimaged in ventral view but also in other viewpoints.

### F. Assessment of Craniofacial Cartilage Dimensions

Use images of Alcian Blue stained tadpoles acquired with a digital camera and stereoscope. Open image analysis software and calibrate the measurement scale. Set the scale for each image before making any measurements. In this protocol we focus on making measurements of elements that can be can be consistently viewed in a ventral view (summarized in Fig. 2 and table 2).

**Table 2:** Summary of cartilages viewed in the protocol.

| Cartilage Element | Anatomical Location | Primary Function |
| --- | --- | --- |
| **Palatoquadrate** | Upper jaw and palate cartilage | Provides structural support for the upper jaw and palate; contributes to mouth width and jaw articulation |
| **Infrarostral** | Anterior midline cartilage at the front of the lower jaw; articulates with Meckel's cartilages | Defines anterior lower jaw shape; supports mouth opening and alignment of the lower jaw |
| **Meckel's Cartilage** | Paired cartilages forming the main anterior lower jaw, extending posteriorly from the infrarostral | Establishes lower jaw length and width; provides scaffold for jaw movement |
| **Ceratohyal** | Paired cartilages posterior to Meckel's cartilages, forming the anterior portion of the hyobranchial apparatus | Supports the floor of the mouth; anchors muscles involved in feeding and buccal pumping |
| **Basihyobranchial** | Midline cartilage within the hyobranchial apparatus, between the ceratohyals | Provides midline structural support; contributes to coordinated feeding and respiratory movements |
| **Ceratobranchials** | Paired elongated cartilages forming the posterior hyobranchial apparatus beneath the cranial base | Define posterior craniofacial structure; support branchial basket and enable feeding and respiratory mechanics |

**Figure 2.**
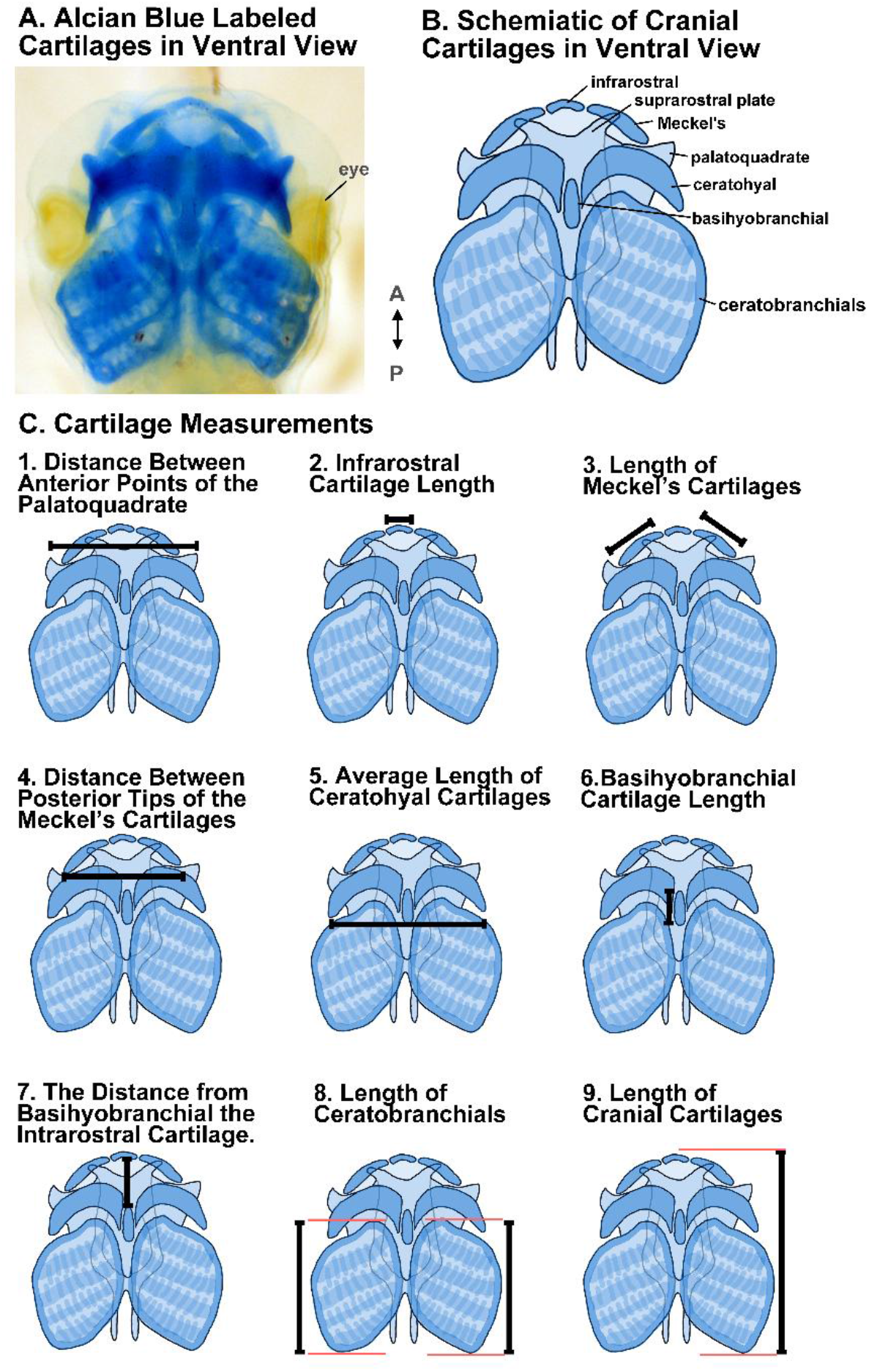
Quantitative analysis of craniofacial cartilages in *Xenopus* tadpoles. (A)Representative ventral view of Alcian Blue stained craniofacial cartilages, highlighting the cartilaginous skeleton used for quantitative analysis. (B) Schematic illustration of craniofacial cartilages in ventral view, with major elements labeled, including the infrarostral plate, suprarostral plate, Meckel’s cartilages, palatoquadrate, ceratohyals, basihyobranchial, and ceratobranchials. Anterior–posterior orientation is indicated. (C) Schematic representations of individual cartilage measurements used for quantitative analysis: (1) distance between the anterior points of the palatoquadrate cartilages, (2) infrarostral cartilage length, (3) length of Meckel’s cartilages, (4) distance between the posterior tips of Meckel’s cartilages, (5) average distance between posterior tips of the ceratohyal cartilages, (6) basihyobranchial cartilage length, (7) distance from the basihyobranchial to the infrarostral cartilage, (8) anterior–posterior length of the ceratobranchial apparatus, and (9) anterior–posterior length of the craniofacial cartilages. Measurement are indicated by black lines while red lines are guide lines.

1. **Distance Between Anterior Points of the Palatoquadrate Cartilage**
  1.1) Identify the paired palatoquadrate cartilages, which lie lateral to Meckel’s cartilages and support the upper jaw.
  1.2) Locate the distal (outermost) tip of the left palatoquadrate cartilage and the distal tip of the right palatoquadrate cartilage.
  1.3) Using the line or distance measurement tool, measure the straight-line distance between these two distal points using the line or distance measurement tool.
  1.4) Record the measurement for each individual tadpole in millimeters (mm).
2. **Infrarostral Cartilage Length**
  2.1) Locate the infrarostral cartilage at the most anterior region of the cranial skeleton.
  2.2) Using the line or distance measurement tool, measure the width of the infrarostral cartilage across its longest dimension from right to left.
  2.3) Record the measurement for each individual tadpole in millimeters (mm).
3. **Length of Meckel’s Cartilages**
  3.1) Identify the left and right Meckel’s cartilages in the ventral view.
  3.2) Using the line or distance measurement tool, measure the length of the left Meckel’s cartilage along its longest axis and record the value in millimeters (mm).
  3.3) Measure the length of the right Meckel’s cartilage in the same manner and record the value in mm.
  3.4) You can also calculate the average of the left and right Meckel’s cartilage lengths for each individual tadpole and record the final value in millimeters (mm).
4. **Distance Between the Posterior Tips of the Meckel’s Cartilages**
  4.1) Locate the left and right Meckel’s cartilages, positioned posterolateral to the infrarostral cartilage.
  4.2) Identify the anterior-most tip of each ceratohyal cartilage.
  4.3) Using the line or distance tool, measure the straight-line distance between these two points.
  4.4) Record the measurement for each individual tadpole in millimeters (mm).
5. **Distance Between Posterior Tips of the Ceratohyal Cartilage**
  5.1) Locate the left and right ceratohyal cartilages, positioned posterolateral to the basihyobranchial cartilage.
  5.2) Identify the posterior-most tip of each ceratohyal cartilage.
  5.3) Using the line or distance measurement tool, measure the straight-line distance between these two points.
  5.4) Record the measurement for each individual tadpole in millimeters (mm).
6. **Basihyobranchial Cartilage Length**
  6.1) Identify the basihyobranchial cartilage at the center of the craniofacial skeleton.
  6.2) Using the line or distance measurement tool, measure the length from anterior to posterior most points.
  6.3) Record the cartilage length for each individual tadpole in square millimeters (mm).
7. **The Distance from the Basihyobranchial to the Infrarostral Cartilage**.
  7.1) Identify the anterior most point of the basihyobranchial cartilage at the center of the craniofacial skeleton and the infrarostral cartilage at the anterior most region of the skeleton.
  7.2) Using the line or distance tool, measure the distance between the two cartilages.
  7.3) Record this distance for each individual tadpole in square millimeters (mm).
8. **The Anterior-Posterior Length of The Ceratobranchial Apparatuses**
  8.1) Draw horizonal guidelines to mark the anterior and posterior extremes of each ceratobranchial apparatus.
  8.2) Measure the distance between the guidelines using the line or distance measurement tool.
  8.3) Record this distance for each the right and left ceratobranchial apparatuses for each individual tadpole in square millimeters (mm).
  8.4) You can also calculate the average of the left and right apparatuses for each individual tadpole and record the value in millimeters (mm).
9. **The Anterior-Posterior Length of Cranial Cartilages**
  9.1) Identify the anterior-most point of the infrarostral cartilage and the posterior-most edge of the ceratobranchial apparatus on the same side.
  9.2) Draw a horizontal guideline at the anterior extreme of the infrarostral cartilage.
  9.3) Draw a second horizontal guideline at the posterior extreme of the ceratobranchial apparatus.
  9.4) Measure the distance between the two guidelines using the line or distance measurement tool.
  9.5) Record this distance for each individual tadpole in millimeters (mm).

### G. Assessment of Craniofacial Cartilage Geometry: Ratios of Dimensions

The following ratios can be taken to provide information about the proportional changes of the cranial cartilages. These can be modified to best quantify specific qualitative observations.

1. **Palatoquadrate to Meckel’s Width Ratio** This ratio reflects the relative width of the upper palate compared to the lower jaw.
  1.1) Use the previously measured palatoquadrate width.
  1.2) Use the previously measured distance between the posterior-most tips of the left and right Meckel’s cartilages.
  1.3) Calculate the ratio by dividing the palatoquadrate distance by the ceratohyal tip distance.
  1.4) Record the calculated ratio for each individual tadpole.
2. **Palatoquadrate to Length of Cranial Cartilages Ratio** This ratio reflects the relative width of the upper jaw/palate compared to overall craniofacial length.
  2.1) Use the previously measured palatoquadrate width.
  2.2) Use the previously measured anterior–posterior length of the cranial cartilages (infrarostral to posterior ceratobranchial edge).
  2.3) Calculate the ratio by dividing the palatoquadrate width by the length of the cranial cartilages.
  2.4) Record the calculated ratio for each individual tadpole.
3. **Meckel’s Width to Infrarostral–Basihyobranchial Distance Ratio** This ratio reflects lower jaw width relative to anterior craniofacial length.
  3.1) Use the previously measured distance between the posterior-most tips of the left and right Meckel’s cartilages.
  3.2) Use the previously measured infrarostral–basihyobranchial distance.
  3.3) Calculate the ratio by dividing the Meckel’s cartilage width by the infrarostral–basihyobranchial distance.
  3.4) Record the calculated ratio for each individual tadpole.
4. **Palatoquadrate to Ceratohyal Width Ratio** This ratio reflects the relative width of the upper jaw/palate compared to the ceratohyal region.
  4.1) Use the previously measured palatoquadrate width.
  4.2) Use the previously measured distance between the left and right ceratohyals.
  4.3) Calculate the ratio by dividing the palatoquadrate width by the ceratohyal width.
  4.4) Record the calculated ratio for each individual tadpole.
5. **Ceratobranchial to Length of Cranial Cartilages Ratio** This ratio reflects the relative length of the posterior branchial region compared to overall craniofacial length.
  5.1) Use the previously measured anterior–posterior length of the ceratobranchial apparatus.
  5.2) Use the previously measured anterior–posterior length of the cranial cartilages.
  5.3) Calculate the ratio by dividing the ceratobranchial length by the cranial cartilage length.
  5.4) Record the calculated ratio for each individual tadpole.
6. **Infrarostral–Basihyobranchial Distance to Length of Cranial Cartilages Ratio** This ratio reflects the contribution of the midline craniofacial axis to overall craniofacial length.
  6.1) Use the previously measured infrarostral–basihyobranchial distance.
  6.2) Use the previously measured anterior–posterior length of the cranial cartilages.
  6.3) Calculate the ratio by dividing the infrarostral–basihyobranchial distance by the cranial cartilage length.
  6.4) Record the calculated ratio for each individual tadpole.

### H. Data Compilation and Statistical Analysis of Measurements and Ratios

1. **Organize and Record Measurements**
  1.1) Create a structured data table with one row per tadpole.
  1.2) Enter all measured craniofacial cartilage dimensions, with each measurement recorded in its own column.
  1.3) Review the dataset to confirm completeness and identify missing or incorrect values.
2. **Calculate Derived Measurements**
  2.1) For paired structures, calculate the average of the left and right measurements for each individual tadpole.
  2.2) Calculate any additional ratios or normalized values in separate columns, ensuring consistent units are used across all entries.
3. **Generate Graphical Summaries**
  3.1) Create comparison plots for each measurement and calculated ratios, displaying Control and Experimental groups side by side.
  3.2) Use consistent labeling, and formatting across all graphs.
  3.3) Display individual data points along with a line indicating the mean or median for each group.
4. **Perform Statistical Testing**
  4.1) For each measurement and calculated ratio, assess whether the data follow a normal distribution using an appropriate normality test.
  4.2) If the data are normally distributed, compare Control and Experimental groups using an unpaired t-test.
  4.3) If the data are not normally distributed, compare Control and Experimental groups using a Mann–Whitney test.
  4.4) Apply a significance threshold of alpha = 0.05 consistently across all comparisons.
  4.5) Record the test used and the resulting p-value for each measured parameter.
5. **Summarize and Interpret Results**
  5.1) Compile a summary table listing group sample sizes, group averages, and p-values for each measurement.
  5.2) Identify measurements that differ significantly between groups.
  5.3) Describe the direction and magnitude of changes using numerical values and graphical trends.

### I. Geometric Morphometric Analysis of Cranial Cartilage Shape

This protocol is a modified version of the protocol previously published for analyzing face shape in *Xenopus laevis* (Kennedy and Dickinson 2014b; Kennedy and Dickinson 2014a).

1. **Landmark placement and coordinate file generation**
  1.1) Select landmarks that capture the geometry of the cranial cartilage skeleton and can be placed consistently in the same relative position across all samples. Landmarks should correspond to discrete, identifiable features of cartilage elements such as cartilage junctions and distal tips (see figure 5A for example). Do not place landmarks on structures that are absent in any sample.
  1.2) Create a figure of your land marks for reproducibility.
  1.3) Open the first image of Alcian Blue stained cranial cartilages and place landmarks using a point-selection tool such as the PointPicker plugin in ImageJ. Adjust landmark positions as needed to ensure precise placement on cartilage boundaries or defined anatomical features. Ensure each landmark is placed in the same order for each sample.
  1.4) Display the landmark coordinate output from ImageJ and copy the coordinates into a spreadsheet program.
  1.5) Leave one empty row above the coordinate data. In column B of this empty row, enter “LM=” followed by the number of landmarks (for example, “LM=15”).
  1.6) In column B of the row immediately below the coordinate data, enter “ID=” followed by a unique identifier for the sample (for example, “ID=C1”).
  1.7) Repeat landmark placement and coordinate export for all tadpoles ensuring that each sample has a unique identifier.
  1.8) Copy the information from columns B and C (which contain the x and y coordinates) into a text document and save the file in plain text (.txt) format. Do not try and save your file as a text document.
2. **Morphometric dataset setup and alignment**
  2.1) Open the morphometric analysis software and create a “New Project”. Assign appropriate names to the project and dataset.
  2.2) Import the landmark coordinate text file as a TPS dataset.
  2.3) Perform a Procrustes fit to align landmark configurations based on cranial cartilage shape. To do this select New Procrustes fit under the Preliminaries tab. Use alignment by principal axis and execute the Procrustes transformation.
  2.4) Generate a covariance matrix from the Procrustes-aligned cartilage landmark data. To do this select Generate Covariance Matrix” under the preliminaries tab and click execute. This will provide correlation values in the results output.
3. **Classifier file creation and import**
  3.1) Create a classifier file in a spreadsheet with two columns labeled “ID” and “TREATMENT.”
  3.2) Enter sample identifiers in the “ID” column and assign the corresponding experimental or control group in the “TREATMENT” column.
  3.3) Save the classifier file as a plain text (.txt) file.
  3.4) Under File, select “Import Classifier Variables” and then choose Match by Identifier. Then select your classifier .txt file.
4. **Discriminant function analysis of cranial cartilage shape**
  4.1) Under the Comparison tab select “Discriminant Function”
  4.2) Select the groups to be compared and run 1,000 permutation tests.
  4.3) Under the Graphics tab you can then visualize shape differences using transformation grids that illustrate changes in cartilage geometry between groups. Landmark displacements represent mean differences in Procrustes-aligned coordinates, while the deformation grid provides a continuous visualization of these changes across the structure. Because deformation is scaled for visualization, interpretation focuses on the spatial pattern of change rather than absolute magnitude.
  4.4) By right clicking on the graph you can adjust vector direction, scale, and grid density as needed to clearly represent cartilage shape differences.
  4.5) Export transformation grids as image files (.jpg).
  4.6) Under the Results tab you can find the Procrustes distance. This is the actual shape difference between group means after aligning all specimens (removing size, position, rotation). It represents the distance between the average shapes of the two groups where 0=identical shapes and higher numbers indicate more difference. There is no universal cutoff and should be interpreted in conjunction with permutation test (p-value) and deformation of the transformation grid.
  4.7) Record the Procrustes distance associated permutation test p-value(s) from the results output. This represents the probability that the observed difference or relationship in your data could occur purely by chance if there is actually no real effect by randomly shuffle group labels. If your p-value is low (p < 0.05), you can say: “The Procrustes distance between groups is significantly greater than expected under random group assignment.”.
5. **Principal component analysis of cranial cartilage shape**
  5.1) Under the Variation tab select Principal Component Analysis (PCA).
  5.2) Navigate to the Graphics tab to visualize a scatter plot.
  5.3) Modify plotted principal components as needed to examine different axes of cartilage shape variation.
  5.4) Right click on the graph to color data points according to treatment group.
  5.5) Export PCA plots as image files (.jpegs).
  5.6) Open the PCA in another image process program and draw ellipses around each colored group.

## Representative Results

To demonstrate the effectiveness of the XenCart Protocol for detecting teratogen-induced craniofacial abnormalities, we conducted a proof of principle experiment using a commercially available e-liquid as the environmental exposure as in published work (Kennedy et al. 2017). *Xenopus* embryos were exposed during a sensitive period in craniofacial development and then later stained with Alcian Blue to visualize cartilage structures.

### A. Size Measurements revealed that e-liquid exposure resulted in smaller cartilage dimensions

Quantitative analysis revealed significant reductions in craniofacial cartilage dimensions following e-liquid exposure, as summarized in Fig. 3 and Table 3. Upper jaw and palatal elements were narrowed, indicating impaired growth or patterning of the upper jaw and primary palate. Lower jaw structures, including the infrarostral and Meckel’s cartilages, were consistently shortened and narrowed. Multiple components of the branchial apparatus, including the ceratohyals, basihyobranchial cartilage, and ceratobranchials, were also reduced, indicating that posterior and midline craniofacial structures were affected in addition to the jaw and palate regions. Together, these findings show that e-liquid exposure causes broad defects across the craniofacial skeleton, with a pattern consistent with a reduction in facial size (Kennedy et al. 2017; Dickinson et al. 2021).

**Table 3:** Summary of Representative Results (A. Size Measurements) and Interpretation.

| # | Measurement | What It Reflects | Fold Change | p-value | Interpretation |
| --- | --- | --- | --- | --- | --- |
| 1 | Palatoquadrate width (anterior points) | Upper jaw and primary palate width | 0.81 | < 0.0001 | Narrowing of the upper jaw region suggests impaired growth or patterning of palatal cartilage consistent with oral clefts. |
| 2 | Infrarostral cartilage length | Anterior midline lower jaw element | 0.63 | < 0.0001 | Strong reduction indicates high sensitivity of anterior midline cartilage reflecting narrowed midface around the mouth. |
| 3 | Meckel's cartilage length | Lower jaw length | 0.76 | < 0.0001 | Shortened lower jaw and may compromise jaw movements |
| 4 | Meckel's cartilage width (posterior tips) | Anterior lower jaw width | 0.83 | < 0.0001 | Reduced anterior lower jaw width suggests narrowing of the anterior face |
| 5 | Ceratohyal cartilage width (posterior tips) | Posterior lower jaw width | 0.91 | < 0.0001 | Mild narrowing indicates posterior lower jaw elements are less affected than anterior elements. |
| 6 | Basihyobranchial cartilage length | Midline hyobranchial support | 0.85 | < 0.0001 | Reduced midline support structure and may compromise feeding-related movements. |
| 7 | Basihyobranchial–infrarostral distance | Anterior–posterior midline craniofacial axis | 0.86 | < 0.0001 | Anterior skeletal shortening in the A-P axis and could reflect defects in mouth structures. |
| 8 | Ceratobranchial apparatus length (A–P) | Posterior craniofacial skeleton | 0.86 | < 0.0001 | Posterior skeletal shortening in the A-P axis and could reflect decreased gill volume. |
| 9 | anterior–posterior length of the craniofacial cartilages | The entire cranial skeleton | 0.67 | < 0.0001 | Total A-P length of cranial cartilages are decreased |

**Figure 3.**
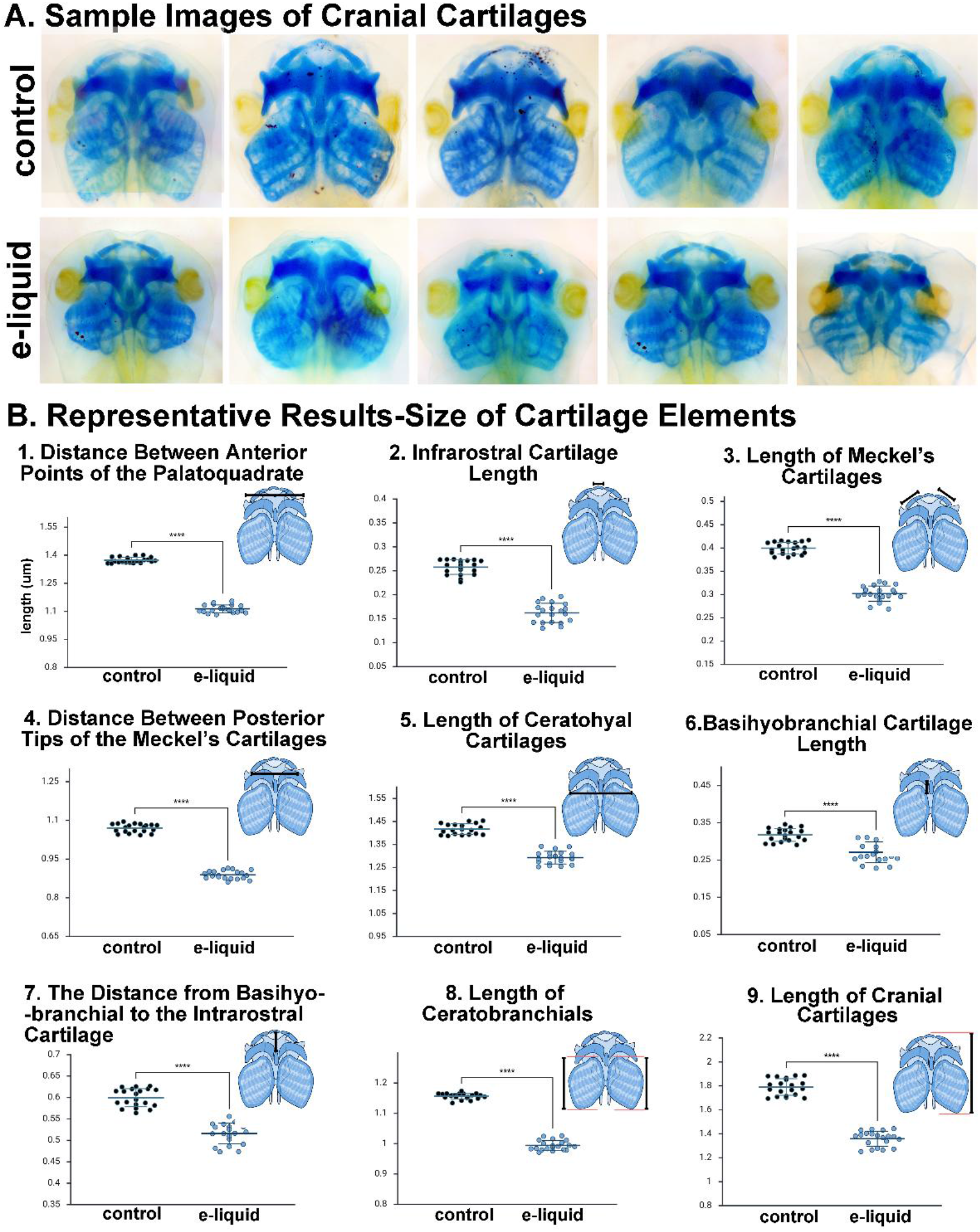
E-liquid exposure disrupts craniofacial cartilage size and proportionality in *Xenopus laevis* tadpoles. (A) Representative Alcian Blue stained cranial cartilage preparations (ventral view) from control and e-liquid–exposed tadpoles. (B) Quantitative analysis of craniofacial cartilage dimensions reveals significant size reductions following e-liquid exposure. Measurements include: (1) distance between the anterior points of the palatoquadrate cartilages, reduced to 0.81-fold of control (Mann–Whitney U test, p < 0.0001); (2) infrarostral cartilage length, reduced to 0.63-fold of control (Mann–Whitney U test, p < 0.001); (3) average length of paired Meckel’s cartilages, reduced to 0.76-fold of control with no detectable left–right asymmetry (unpaired t test, p < 0.0001); (4) distance between posterior tips of the Meckel’s cartilages, reduced to 0.83-fold of control (unpaired t test, p < 0.0001); (5) distance between posterior tips of the ceratohyal cartilages, reduced to 0.91-fold of control (unpaired t test, p < 0.0001); (6) basihyobranchial cartilage length, reduced to 0.85-fold of control (unpaired t test, p < 0.0001); (7) distance from the basihyobranchial to the infrarostral cartilage, reduced to 0.86 -fold of control (Mann–Whitney U test, p < 0.0001); and (8) anterior–posterior length of the ceratobranchial apparatus, reduced to 0.86-fold of control (unpaired t test, p < 0.0001). (9) anterior–posterior length of the cranial cartilages, reduced to 0.67-fold of control (unpaired t test, p < 0.0001)Individual data points represent single embryos, with bars indicating mean ± standard deviation.

### B. Ratios of dimensions revealed that e-liquid exposure altered changed the proportional relationships between cranial cartilages

Analysis of craniofacial cartilage ratios revealed alterations in proportional relationships following e-liquid exposure (summarized in Fig. 4 and table 4). Ratios comparing upper jaw width to lower jaw width and to overall craniofacial length were modestly but significantly reduced, indicating altered scaling of the palatoquadrate relative to anterior facial structures. Lower jaw width relative to anterior craniofacial length was also reduced, further supporting disproportionate changes within the anterior face. The ratio of palatoquadrate to ceratohyal width was increased, suggesting region-specific differences in scaling between upper jaw and hyobranchial elements. Ratios assessing the contribution of posterior and midline structures to total craniofacial length, including ceratobranchial length and the infrarostral–basihyobranchial axis, were reduced, indicating a relative shortening of posterior and midline craniofacial components.

**Table 4:** Summary of Representative Results (B. Ratios of Dimensions) and Interpretation.

| # | Measurement | What It Reflects | Fold Change | p Value | Interpretation |
| --- | --- | --- | --- | --- | --- |
| 1 | Palatoquadrate: Meckel's width ratio | Upper jaw width relative to lower jaw width | 0.98 | 0.0004 | Subtle narrowing of the upper palate relative to the lower jaw. |
| 2 | Palatoquadrate: cranial cartilage length ratio | Upper jaw width relative to overall craniofacial length | 0.98 | < 0.0001 | Upper jaw width does not scale normally with craniofacial length consistent with a narrower head. |
| 3 | Meckel's width: infrarostral–basihyobranchial distance ratio | Lower jaw width relative to anterior craniofacial length | 0.97 | 0.018 | Altered proportional relationships within the anterior craniofacial region. |
| 4 | Palatoquadrate: ceratohyal width ratio | Upper palate relative to ceratohyal region | 1.13 | 0.0004 | Relative expansion of palatoquadrate compared to ceratohyals suggests region-specific scaling differences. |
| 5 | Ceratobranchial: cranial cartilage length ratio | Posterior branchial region relative to total craniofacial length | 0.88 | < 0.0001 | Reduced contribution of posterior branchial structures to overall craniofacial length. |
| 6 | Infrarostral–basihyobranchial: cranial cartilage length ratio | Midline craniofacial axis relative to total craniofacial length | 0.88 | < 0.0001 | Disproportionate shortening of the midline craniofacial axis. |

**Figure 4.**
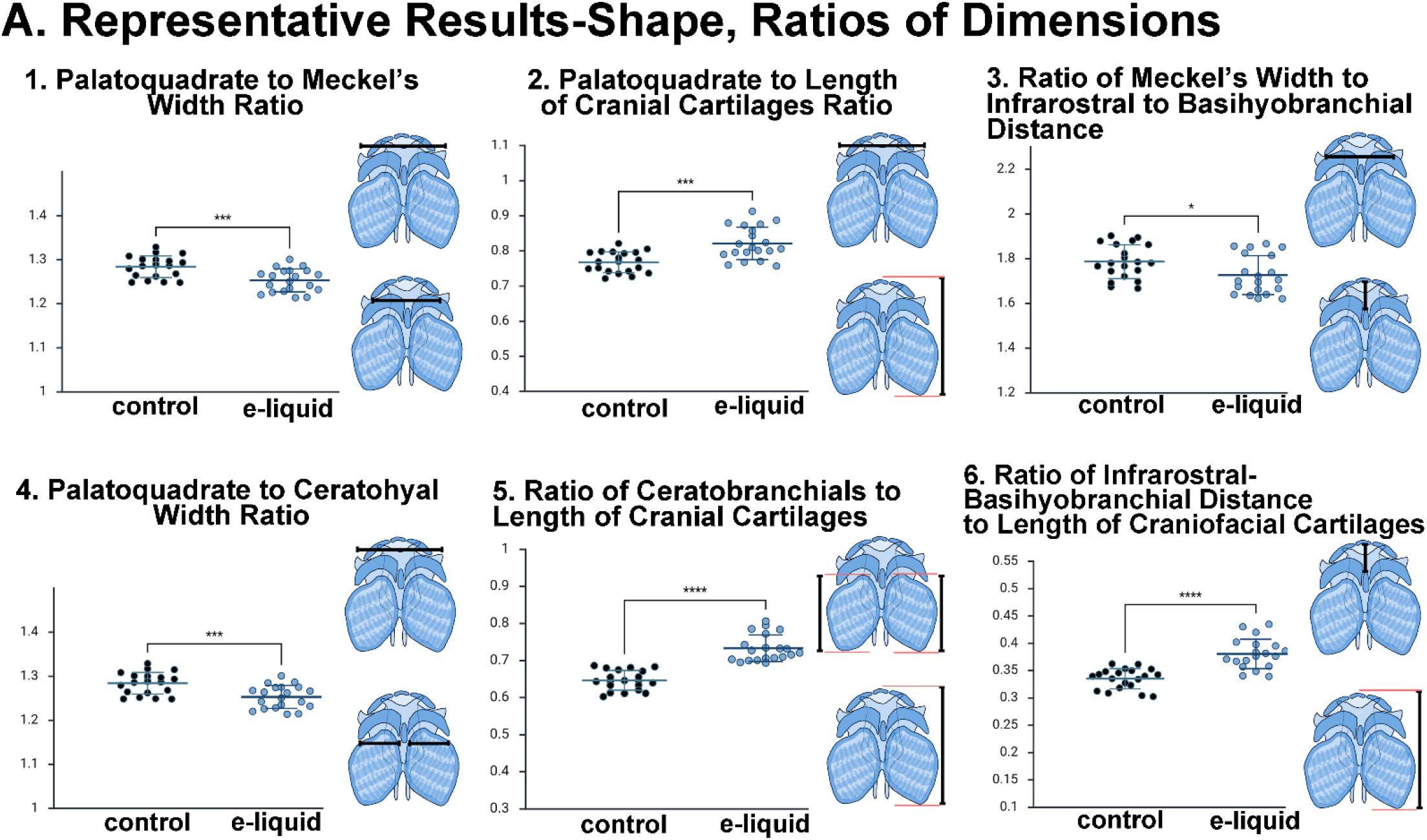
Ratio-based analysis reveals region-specific changes in craniofacial cartilage geometry following e-liquid exposure. (A) Representative scatter plots showing ratios of craniofacial cartilage dimensions in control and e-liquid–exposed *Xenopus* tadpoles. Each point represents an individual embryo; bars indicate mean ± SEM. Schematic cartoons illustrate the measurements used for each ratio. (1) Palatoquadrate to Meckel’s width ratio, reflecting relative upper palate width compared to the lower jaw, was modestly but significantly reduced in e-liquid–exposed tadpoles [0.98-fold decrease; t test, p = 0.0004]. (2) Palatoquadrate to cranial cartilage length ratio, reflecting upper palate width relative to overall craniofacial length, was significantly reduced [0.98-fold decrease; t test, p < 0.0001]. (3) Meckel’s width to infrarostral–basihyobranchial distance ratio, reflecting lower jaw width relative to anterior craniofacial length, was significantly reduced [0.97 -fold decrease; Mann–Whitney U test, p = 0.018]. (4) Palatoquadrate to ceratohyal width ratio, reflecting relative upper palate width compared to the ceratohyal region, was significantly increased [1.13-fold increase; t test, p = 0.0004]. (5) Ceratobranchial to cranial cartilage length ratio, reflecting the contribution of the posterior branchial region to overall craniofacial length, was significantly reduced [0.88-fold decrease; Mann–Whitney U test, p < 0.0001]. (6) Infrarostral–basihyobranchial distance to cranial cartilage length ratio, reflecting the contribution of the midline craniofacial axis to total craniofacial length, was significantly reduced [0.88-fold decrease; t test, p < 0.0001].

**Figure 5.**
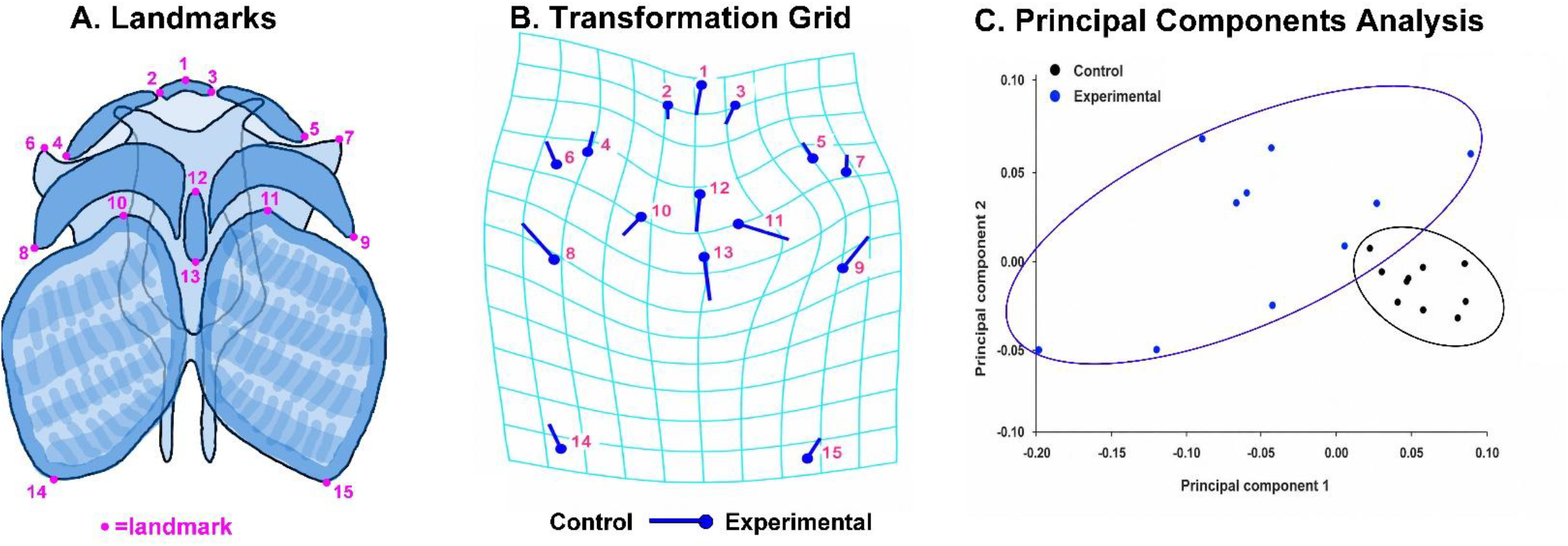
Representative geometric morphometric analysis of cranial cartilage shape. (A) Schematic showing the craniofacial cartilage elements and the 15 anatomical landmarks used for geometric morphometric analysis. Landmarks were placed at conserved positions across specimens to capture overall craniofacial shape. (B) Discriminant function analysis (DFA) visualized as a transformation grid, showing the shape changes that distinguish control and experimental groups. Blue vectors indicate the direction and relative magnitude of landmark displacement from the control mean shape toward the experimental mean shape. Grid deformation illustrates the corresponding changes in overall craniofacial shape. Procrustes distance = 0.0774 (p = 0.005). (C) Principal components analysis (PCA) of landmark coordinates showing the distribution of individual specimens in morphospace. Each point represents one specimen, and ellipses summarize group-level variation. Separation between control and experimental samples along principal component 1 indicates treatment-associated differences in craniofacial shape.

### C. Morphometric analysis revealed that e-liquid exposure caused overall shape change in the cranial cartilage

Geometric morphometric analysis indicated that e-liquid exposure caused changes in the shape of the cartilages in the head of the *Xenopus* tadpole (summarized in Fig. 5). Discriminant function analysis indicated a Procrustes distance of d = 0.0774 that was significantly greater than expected under permutation of group labels (p = 0.005), indicating a non-random shape difference between groups. Transformation grids illustrate the pattern of shape change between group means. Landmark displacement vectors were consistent with an overall narrowing and anteroposterior compression of the craniofacial skeleton.

Principal component analysis showed that exposed specimens were shifted towards negative PC1 values whereas control embryos clustered predominantly in the positive values. This separation along the dominant axis of shape variation indicated that e-liquid exposure altered mean cartilage shape. In addition, e-liquid exposed embryos were also more broadly placed along the PC1 axis indicating shape variability. This could suggest variable sensitivity among individual embryos to the teratogenic effects of the e-liquid.

In summary our quantitative analysis of craniofacial cartilages revealed reproducible morphological differences between treated and control tadpoles, confirming that this workflow can sensitively detect changes in cartilage patterning. These findings validate the protocol as a reliable approach for assessing how emerging environmental toxicants alter craniofacial development.

## Discussion

Most craniofacial cartilage analyses in *Xenopus* have been qualitative and focused on identifying affected elements rather than quantifying changes in the spatial relationships and geometry of the craniofacial skeleton that ultimately shape function and structure. Because facial form and function depend on precise relationships among skeletal size, shape, and position, quantitative assessment of craniofacial geometry is essential for moving beyond descriptive phenotypes. To address this need we have established XenCart, a simple, flexible Alcian Blue staining and measurement framework that enables standardized, quantitative assessment of craniofacial cartilage size and geometry. The protocol which builds on established vertebrate staining approaches while incorporating timing flexibility that makes it suitable for both research and instructional settings, including course-based undergraduate research experiences.

As a proof-of-principle, we show how our protocol revealed coordinated changes across the craniofacial skeleton following an environmental exposure (e-liquid), linking external facial phenotypes identified in previous work (Kennedy et al. 2017; Dickinson et al. 2021)to underlying skeletal alterations. Importantly, several proportional measurements and geometric morphometrics were altered independently of overall size, indicating that e-liquid exposure affects craniofacial geometry in addition to growth. The representative results were generated from approximately 10–20 embryos pooled from three biological replicates; future studies should have increased samples as well as preserve replicate identity to compare both within and across biological replicates to better assess variability and reproducibility. Regardless, these findings point to disruption of region-specific developmental programs that govern craniofacial patterning or growth.

### Limitations and Future Directions

Although the XenCart protocol provides an adaptable and practical approach for quantifying craniofacial cartilage morphology, several limitations should be considered. The current workflow focuses primarily on ventral views and on cartilage elements that can be consistently visualized from this orientation, which limits assessment of dorsal structures and how cartilage dimensions or geometry may change when viewed laterally or dorsally. In addition, the present analysis is based on 2D images, which can detect changes in size, proportion, and shape but cannot fully capture depth, curvature, volume, or three-dimensional spatial relationships among cartilage elements. Future work should expand the protocol to include standardized dorsal and lateral imaging, as well as 3D approaches to quantify cartilage geometry more completely. Our immediate goals are to also use the standardized images, landmark coordinates, and quantitative measurements generated through this workflow to train AI-based image analysis models for faster, more automated, and potentially more sensitive detection of craniofacial cartilage defects. These future directions will allow XenCart to develop from a simple 2D screening method into a more sophisticated and scalable platform for analyzing craniofacial cartilage structure and developmental perturbation in Xenopus.

In conclusion, this study presents a framework for quantitative analysis of craniofacial cartilage in *Xenopus* larvae that is well suited for both research and instructional settings. The XenCart protocol provides a broadly applicable and accessible platform for assessing how genetic, environmental, and pharmacological perturbations alter craniofacial development and function.

## Acknowledgements

This protocol was developed and data generated in independent undergrad projects and two CURE courses taught by Dr. Dickinson in 2025 in the School of Life Sciences and Sustainability at Virginia Commonwealth University. Students included, Leona Bhandari, Annmarie Gendy, Vinh Hoang, Nichole Holcombe, Riya Maurya, Eva Smart, Jadyn Williams, Bedria Abdulmelik, Umar Aziz, Kriti Khadka, Layla Lawal, Aliya Mulla, Viet Nguyen, Seyram Nkulenu, Jared Reyes, Deja Smith, Skyler Yann, Joseph Yao and Claudia Lizama.

